# Sleep-dependent structural neuroplasticity after a spatial navigation task: A diffusion imaging study

**DOI:** 10.1101/2022.07.05.498806

**Authors:** Thomas Villemonteix, Michele Guerreri, Michele Deantoni, Evelyne Balteau, Christina Schmidt, Whitney Stee, Hui Zhang, Philippe Peigneux

**Affiliations:** UR2NF-Neuropsychology and Functional Neuroimaging Research Unit affiliated at CRCN – Centre for Research in Cognition and Neurosciences and UNI - ULB Neuroscience Institute, Université Libre de Bruxelles (ULB), Brussels, Belgium; Laboratoire Psychopathologie et Processus de Changement, EA2027, Paris 8 University, Rue de la Liberté 2, 93526 Saint-Denis, France; Department of Computer Science & Centre for Medical Image Computing, University College London, UK3; Sleep & Chronobiology Group, GIGA-CRC-In Vivo Imaging Research Unit, University of Liège, Liège, Belgium

## Abstract

Evidence for sleep-dependent changes in micro-structural neuroplasticity remains scarce, despite the fact that it is a mandatory correlate of the reorganization of learning-related functional networks. We investigated the effects of post-training sleep on structural neuroplasticity markers measuring standard diffusion tensor imaging (DTI) mean diffusivity (MD) and the revised biophysical neurite orientation dispersion and density imaging (NODDI) free water fraction (FWF) and neurite density (NDI) parameters that enable disentangling whether MD changes result from modifications in neurites or in other cellular components (e.g., glial cells). Thirty-four healthy young adults were scanned using diffusion weighted imaging [DWI] on Day1 before and after 40-minutes route learning (navigation) in a virtual environment, then were sleep deprived (SD) or slept normally (RS) for the night. After recovery sleep for 2 nights, they were scanned again (Day4) before and after 40-minutes route learning (navigation) in an extended environment. Sleep-related microstructural changes were computed on DTI (MD) and NODDI (NDI and FWF) parameters in the cortical ribbon and subcortical hippocampal and striatal regions of interest (ROIs). Results disclosed navigation learning-related decreased DWI parameters in the cortical ribbon (MD, FWF) and subcortical (MD, FWF, NDI) areas. Post-learning sleep-related changes were found at Day4 in the extended learning session (pre- to post-relearning percentage changes), suggesting a rapid sleep-related remodelling of neurites and glial cells subtending learning and memory processes in basal ganglia and hippocampal structures.

## 1. Introduction

Neuroplasticity is the process by which our brain is able to structurally and functionally remodel as a result of cognitive experience (Assaf, 2018). Neuroplasticity mechanisms supporting learning and memory consolidation are operant both online, during the acquisition of novel information, and offline over two timescales after the actual learning episode (Stee & Peigneux, 2021). On the short term is the process initiated during learning to write and stabilize memory traces within local circuits, ranging over a timescale of seconds to hours. At a longer term takes place the processes that reorganizes memory traces at the systems-level, on a timescale of hours to several weeks (Wamsley, 2019). Studies from cell recording in rodents to behavioural and functional neuroimaging data in human indicate that post-learning sleep plays a key contributing role in the systems-level consolidation of recently acquired information, although offline memory consolidation can also take place during states of wakefulness (Peigneux, 2015; Rasch & Born, 2013; Wamsley, 2019). Nowadays, evidence for sleep-dependent changes in micro-structural neuroplasticity remains scarce, despite the fact that it is a mandatory correlate of the reorganization of functional learning-related networks (Stee & Peigneux, 2021).

Among key cognitive systems, spatial memory is responsible for recording information about one’s environment and spatial orientation (Do et al., 2021). Spatial memory shares similar neuroanatomical foundations in human beings and animals (Burgess et al., 2002), which makes spatial learning an attractive paradigm to study the effects of sleep, with unique opportunities for translational inferences (Peigneux et al., 2004). Functional brain imaging studies typically found spatial navigation associated with activity in a distributed cerebral network including the hippocampus, dorsal striatum and entorhinal, parahippocampal, retrosplenial and frontal cortices (Epstein et al., 2017). Hippocampus and entorhinal cortex are proposed to subtend the creation of a map-like spatial code, i.e., a representation of the structure of the environment. Parahippocampal and retrosplenial cortices are thought to provide critical inputs anchoring the cognitive map to fixed environmental landmarks (Epstein et al., 2017; Gahnstrom & Spiers, 2020). Spatial coding is not limited to medial temporal structures; spatial representations are also associated with orbito-frontal (Wikenheiser et al., 2021) and/or parietal (Brodt et al., 2018; Schott et al., 2019) cortex activity. Besides map-based representations, spatial navigation can also rely on stimulus–response associations supporting the development of habit-based route strategies. Human functional Magnetic Resonance Imaging (fMRI) and animal lesion studies evidenced a role for the dorsal striatum in the formation of habits and habit-based navigation (Orban et al., 2006; Rauchs, Orban, Balteau, et al., 2008; Wunderlich et al., 2012). The dorsal striatum plays a complementary role during flexible navigation, through the implementation of reinforcement learning computations supporting encoding of transitions structure and forward planning in the caudate nuclei (Gahnstrom & Spiers, 2020).

The two-stage theory of systems-level memory consolidation proposes that declarative and spatial memory traces are initially encoded into the hippocampus, which serves as a transient storage site, and are then gradually transferred to the neocortex for long-term storage. Sleep is thought to play a key role in this process, thanks to the oscillatory rhythms occurring during deep non-rapid eye movement (NREM) sleep (Binder et al., 2019; Born & Wilhelm, 2012; Buzsáki, 1998), namely the orchestrated interplay between neocortical slow oscillations, thalamo-cortical spindles and hippocampal sharp-wave ripples. Rodent studies showed the spontaneous replay of learning-related neural patterns of hippocampal place cell activity (e.g., Ólafsdóttir et al., 2018). Accordingly, in human, increased activity was found during NREM sleep in navigation-related hippocampal and parahippocampal regions in participants trained to explore a virtual city before sleep, as compared to subjects trained on another, non-hippocampus-dependent procedural memory task (Peigneux et al., 2004). Besides, fMRI and magnetoencephalography (MEG) studies provided evidence that post-learning sleep leads to a reorganization of the brain patterns underlying the retrieval of topographical or declarative memories (Orban et al., 2006; Rauchs, Orban, Schmidt, et al., 2008; Urbain et al., 2016). Indeed, functional MRI studies showed that navigation in a recently learned environment, initially subtended by hippocampus-based spatial strategies, becomes more contingent on striatal activity a few days later when participants are allowed to sleep on the first night after learning (Orban et al., 2006; Rauchs, Orban, Schmidt, et al., 2008), suggesting that sleep favoured a shift toward a habit-based, cognitively less demanding navigation strategy.

Memory-related neuroplasticity mechanisms have been mostly investigated in humans measuring functional brain responses (e.g, BOLD responses with fMRI, regional cerebral blood flow (rCBF) with positron emission tomography (PET), electroencephalography (EEG)/MEG signal …), but the underlying micro-structural brain changes are more elusive to detect at limited time scales such as hours or even days. In this respect, advances in diffusion magnetic resonance imaging (dMRI) techniques have opened new avenues to explore the (micro)structural neuroplasticity changes supporting memory processes in the human brain (Assaf, 2018). Diffusion tensor imaging (DTI) studies showed that mean diffusivity (MD), a global marker of tissue microstructure, can track the microstructural changes associated with learning on the short term. Investigating spatial learning-related neuroplasticity, Sagi et al. (2012) evidenced decreased mean diffusivity (MD) in the grey matter bilaterally in the hippocampus and parahippocampus in participants repeatedly trained on the same route for 120 minutes in a video game task, as compared to control participants driving random routes A similar study evidenced decreased MD in hippocampus, precuneus, superior frontal gyrus, left insula, supramarginal, superior temporal and fusiform gyri following a 60-minutes training session in which participants drove 20 times the same route (i.e., learning it), as compared to a control group who practiced driving 20 different routes (Keller & Just, 2016). To the best of our knowledge, only one study tested an effect of sleep on micro-structural plasticity; results showed decreased mean cortical MD after 12 hours of intense training on a driving simulation game and executive function tasks, an effect magnified after 24 h of task practice combined with sleep deprivation, that reverted after recovery sleep (Bernardi et al., 2016).

An inherent limitation of DTI is that even if sensitive to tissue’s microstructure, it merely provides a composite view of all cellular components within the measured voxel and does not provide information on its internal partitions. Tavor et al. (2013) evidenced short-term plasticity changes using both a conventional DTI approach and the biophysical composite hindered and restricted diffusion (CHARMED) approach, that models the tissues in a voxel as the integration of multiple components (i.e., hindered and restricted compartments) and provides an estimate for each of them (Sagi et al., 2012). In this study, decreased MD was found again after 120 minutes car-racing in the left hippocampus, bilateral para-hippocampal gyrus and bilateral insula. MD changes were paralleled by an increased water volume in the restricted compartment (Fr) of the CHARMED model, tentatively proposed to reflect the remodelling of dendrite and glial processes in the grey matter (GM) (Tavor et al., 2013). Furthermore, increased Fr was detected in hippocampal and parahippocampal areas already after 45 minutes of learning, while DTI failed to reflect such changes, suggesting CHARMED a more sensitive method to study structural plasticity. However, CHARMED was mostly designed for white matter (WM) modelling and is thus less suited for describing GM changes. An alternative to CHARMED is the neurite orientation dispersion and density imaging (NODDI) model, a biophysical model specifically designed to describe both WM and GM (Zhang et al., 2012). The NODDI model assumes that water molecules in neuronal tissue can be considered within three separate compartments: (a) free water, representing CSF; (b) restricted water representing neurites; and (c) hindered water, representing diffusion within glial cells, neuronal cell bodies and the extracellular environment. This enables estimation of parameters including the free water fraction index (FWF), the neurite density index (NDI) and the orientation dispersion index (ODI) that provide complementary information regarding the extent of neurite orientation variability in the space. Hence, compared to DTI-derived metrics, NODDI indices have the potential to provide a more specific characterization of the microstructural changes related to neuroplasticity.

As mentioned above, evidence for sleep-dependent changes in micro-structural neuroplasticity remains scarce (Stee & Peigneux, 2021). In the present dMRI study, we investigated the effects of a post-training sleep manipulation on structural markers of neuroplasticity. Participants were scanned using diffusion weighted imaging (DWI) on Day 1 before and after a 40-minutes route learning (navigation) session in a virtual environment (Peigneux et al., 2004, 2006), then sleep deprived (SD) for the whole night or allowed to sleep normally (RS). They all had then 2 nights with normal sleep. On Day 4, they were scanned again using DWI before and after a 40-minutes route learning (navigation) session in an extended version of the virtual environment. We used dMRI derived metrics from both DTI and NODDI approaches to assess microstructural changes in the cortical ribbon and in subcortical regions of interest (ROIs). Results disclosed learning-related decreased parameters in the cortical ribbon and subcortical areas, and post-learning sleep-related changes in the extended learning session at Day 4 in ROIs caudate nucleus and hippocampus, suggesting an early sleep-related remodelling of both neurites and glial processes in subcortical regions.

## 2. Material and Methods

### 2.1. Participants and Procedure

34 healthy volunteers gave written informed consent to participate in this study approved by the Ethics Committee of the University of Liege, Belgium (reference B70720156274). They were paid for their participation. All procedures were performed following relevant guidelines and regulations. One participant was excluded due to abnormal structural MRI scan, and one participant due to a corrupted data file in one of the diffusion imaging acquisitions. The final sample consisted of 32 right-handed (Laterality Quotient Score > 70 (Oldfield, 1971) participants (17 males; mean age, 23.5 years; range, 19–31 years). Self-reported questionnaires assessed sleep habits (PSQI (Buysse et al., 1989)) and circadian topology (MEQ (Horne & Ostberg, 1976)) for the past month before inclusion. Exclusion criteria were extreme chronotype (MEQ score > 69 or < 31) and bad sleep quality (PSQI score > 5), prior history of neurological or psychiatric disorder, or consumption of psychotropic drugs.

The experimental timeline is illustrated Figure 1. On Day 1, sessions were conducted in the afternoon from 13:00 to 20:00. On that day, mean wake-up time as assessed by self-reported questionnaires was 08:43 (SD = 44 minutes). Participants were first scanned with a diffusion-weighted imaging session (DWI-1), then had four successive 8-minutes blocks outside of the scanner to explore and learn the Level 1 of a virtual city (see below). They were then scanned with functional magnetic resonance imaging (fMRI) while performing a navigation test where they had to reach as fast as possible 10 targets within the learned environment (Day 1 Navigation Performance Score), and then scanned again in a second diffusion imaging session (DWI-2). Participants were then randomly assigned either to the regular sleep (RS; n = 15; 8 males) or to the sleep deprivation (SD; n = 17; 9 males) condition. Participants in the RS group were allowed to sleep as usual at home and requested to follow regular sleep habits for 3 experimental nights, based on their usual schedule. In the SD group, participants were kept awake in the laboratory during the first post-learning night. During this SD night, their physical activity was maintained as low as possible under constant supervision by the experimenters. Participants remained most of the time in a seated position, and were allowed to read, chat, play quiet games and watch movies. They received isocaloric meals at regular intervals, and water ad libitum. At 8:00 am, they were allowed to leave the lab, and were instructed to have non-dangerous daytime activities and abstain as much as possible from napping until bedtime. They then slept normally at home and were requested to follow regular sleep habits the two following nights. Sleep quality and quantity for the nights from before the first learning session (Day 1) to the last testing session (Day 4) was assessed using a daily standardized questionnaire (Ellis et al., 1981), controlled by visual inspection of continuous actimetric recordings (ActiGraph, wGT3X-BT Monitor, USA). On Day 4, SD and RS participants came back to the laboratory for a second experimental session in the afternoon, at the same time than at Day 1. First, they were scanned with diffusion imaging (DWI-3). Afterwards, they were scanned with functional magnetic resonance imaging (fMRI) while performing a navigation test where they had to reach as fast as possible 10 targets within the learned L1 environment (Day 4 Navigation Performance Score). They were then trained outside of the MRI scanner for an additional four 8-minutes blocks on an extended version of the navigation task (Level 1 plus Level 2 of the town; Figure 1), in which the targets and environments of the initial training were incorporated. They were then scanned again with the diffusion sequence (DWI-4) and finally tested on the extended version of the navigation task outside of the MRI environment (Day 4 Relearning score). A high-resolution structural MRI scan was also acquired for each participant at the end of the brain imaging session (Day 1 or Day 4). In this protocol, 5-minutes long resting-state fMRI sessions were also acquired after each DWI session. Resting-state fMRI results have been reported in a previous publication (Deantoni et al., 2021) and will not be discussed here.

**Figure 1.**
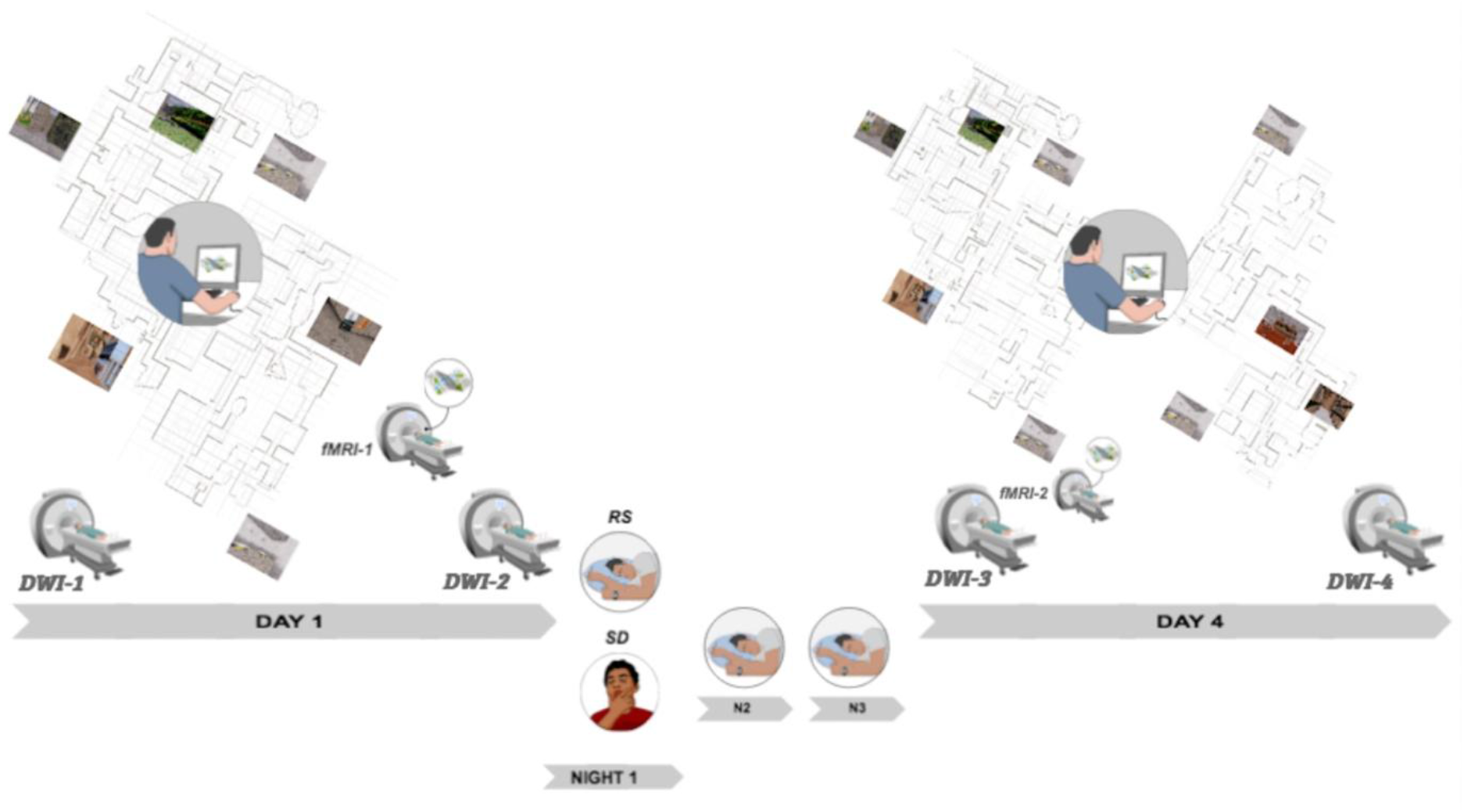
Experimental Protocol. On Day 1, participants are scanned with diffusion-weighted imaging before (DWI-1) and after (DWI-2) out-of-scanner learning of Level 1 of the virtual town. They are also tested for navigation performance in a fMRI session (fMRI-1) at the end of learning before DWI-2. Participants then either sleep normally (RS) or are sleep deprived (SD) during post-learning Night 1, and then sleep normally on Nights 2 and 3. On Day 4, RS and SD participants are scanned again with diffusion (DWI-3) then tested again for navigation performance in a fMRI session (fMRI-2). Afterward, they are trained on an extended version (both Level 1 and 2) of the virtual town and then scanned a last time (DWI-4). Finally, participants are tested on the extended version of the virtual town outside of the MRI environment (Retrieval extended, not illustrated). Top left and right panels provide an aerial representation of the town map (not seen by participants who navigated from within the environment, see sample pictures) with Level 1 (left, day 1) and 2 (right, day 4) environments.

### 2.2. Navigation task

The navigation task is described in detail elsewhere (Deantoni et al., 2021), only essential information is reported here. Participants were trained in a previously used virtual navigation environment (Orban et al., 2006; Peigneux et al., 2006; Rauchs, Orban, Balteau, et al., 2008), that features a complex town made of two levels communicating through two teleports (Figure 1). Each level is composed of three districts, distinct from each other both visually (architecture, objects …) and auditorily (background sounds and music) and containing each one target location. Participants are explicitly instructed to learn their way in the city by moving freely within the environment. At both Day 1 and Day 4 learning sessions, participants are exposed to a test trial at the end of each 8-minute exploration block, i.e., they are assigned a starting point and instructed to reach a given target location as fast as possible in a maximal time of 28 s (Day 1) or 60 s (Day 4), i.e., the shortest possible duration from starting point to target destination within L1 (Day 1) or from L1 to L2 (Day 4). At the end of the four 8-minute blocks of free exploration, participants are exposed to 10 test trials in a row to estimate their navigation performance. A quantitative performance measure is computed for each trial as the shortest distance remaining between the subject’s actual location at the end of the trial time limit and the location of the target destination, based on video screen recordings of their navigation (Deantoni et al., 2021; Orban et al., 2006; Peigneux et al., 2006; Rauchs, Orban, Balteau, et al., 2008). Participants were also tested on 10 L1 navigation trials during a fMRI session on Day 1 after learning before DWI-2 and on Day 4 before relearning after DWI-3.

### 2.3. Diffusion Imaging

#### 2.3.1. Data Acquisition

Brain MRI data were acquired on a whole-body 3T scanner (Magnetom Prisma, Siemens Medical Solutions, Erlangen, Germany). Diffusion MRI images were acquired with a 64-channel head coil, using a multislice 2D single-shot spin-echo echo-planar imaging with monopolar gradients. Parallel imaging with the iPAT technique was used with a reduction factor of 2, one excitation, and 2mm interleaved axial slices with no gap at an isotropic resolution of 2x2 mm on a 96x96 matrix. The echo time (TE) and repetition time (TR) were 69ms and 7400ms. There were 70 slices (b-values: 650, 1000 and 2000 sec/mm2 with 15, 30 and 60 directions respectively; 13 additional interleaved brain volumes were also acquired with no diffusion weighting, and two extra b=0 volumes were acquired with opposite phase encoding for susceptibility induced distortion correction). For anatomical reference, a high-resolution T1-weighted structural image was acquired for each subject (T1-weighted 3D magnetization-prepared rapid gradient echo (MPRAGE) sequence, TR = 1900 ms, TE = 2.19 ms, FA = 9 deg, inversion time (TI) = 900 ms, FoV = 256 × 240 mm2, matrix size = 256 × 240 × 224, voxel size = 1 × 1 × 1 mm3, acceleration factor in phase-encoding direction R = 2).

For task-based fMRI, multi-slice T2*-weighted functional images were acquired with a gradient-echo echo-planar imaging EPI) sequence using axial slice orientation and covering the whole brain (36 slices, FoV = 216 × 216 mm2, voxel size 3 × 3 × 3 mm3, 25% interslice gap, matrix size = 72 × 72 × 36, repetition time (TR) = 2260 ms, echo time (TE) = 30 ms, flip angle (FA) = 90 deg). Processing and analyses of task-based fMRI data are reported in Deantoni et al. (2021).

#### 2.3.2. MRI data processing

In this section we detail the steps towards the estimation of the diffusion MRI (dMRI) derived metrics as well as towards the determination of correspondence across subjects and across timepoints. Microstructural changes associated to fast learning episodes have been previously reported in subcortical as well as in neocortical areas (Brodt et al., 2018; Keller & Just, 2016; Sagi et al., 2012), hence our analysis included both. We derived two streams of analysis optimized for each of them. Microstructural changes in the cortex were estimated via a surface-based analysis by projecting the dMRI metrics on the cortical surface. Microstructural changes in the grey matter of the hippocampus, caudate nucleus and putamen were assessed via a region of interest (ROI)-based analysis. In both cases we leveraged the anatomical information from T1-weighted (T1-w) images to establish inter-subject correspondence. Intra-subject group correspondence between different timepoints was established using the information from DWI.

##### 2.3.2.1 Pre-processing

Correspondence across subjects, for both the surfaced-based and the ROI-based analysis, was determined via the anatomical (T1w) images via FreeSurfer software (https://surfer.nmr.mgh.harvard.edu/, v7.1.1). To improve the anatomical image analysis robustness, pre-processing steps inspired by the minimal pre-processing pipeline for the Human Connectome Project (Glasser et al., 2013) were applied prior to the FreeSurfer analysis. First, anatomical images were reoriented to match the orientation of the standard MNI152 template images. Next, they were cropped to a smaller FOV removing the neck and the lower part of the head, and finally, they were aligned to match the MNI152 template via the anterior to posterior commissure (“ACPC”) alignment procedure (Glasser et al., 2013). These steps were implemented via the FMRIB Software Library (FSL, v6.0) (Jenkinson et al., 2012; Smith et al., 2004; Woolrich et al., 2009). A preliminary analysis showed that the anatomical image marked bias field affected the FreeSurfer analysis outcome quality, hence all images underwent an extra bias field correction step via the N4ITK algorithm (Tustison et al., 2010). Finally, in order to assist the final registrations to MNI space and to assist FreeSurfer with its brain mask creation process, a rough brain mask was generated using the method described in Glasser et al (2013).

For dMRI, the pre-processing steps included susceptibility induced distortion correction, eddy current induced distortion correction and head motion correction. Susceptibility induced distortions were estimated from b=0 volumes with opposed phase encoding via FSL’s topup (Andersson et al., 2003). FSL’s eddy routine was used to simultaneously correct for susceptibility and eddy current induced distortions, as well as for head motion (Andersson & Sotiropoulos, 2016).

##### 2.3.2.2 diffusion MRI (dMRI) models

As a reminder, NODDI models brain microarchitecture using three complementary components within a voxel volume: (1) intra-neurite (modelling the space bounded by the neurite membranes and myelin sheaths), (2) extra-neurite (modelling the space outside of neurites, including glial cells and somas), and (3) free water (accounting for the presence of cerebrospinal fluid or oedema in the tissues) (Zhang et al., 2012). NODDI’s output is a set of indices aimed at describing neurite morphology. The neurite density index (NDI) accounts for the intra-neurite compartment signal fraction. The free-water fraction (FWF) is the free-water compartment signal fraction. The orientation dispersion index (ODI) measures the directional variability of neurites. In this work we used a revised version of NODDI which addresses some of the original technique limitations when extended to GM (Guerreri et al., 2018, 2020). NODDI revised version has been shown to be more sensitive to subtle microstructural changes in the cortex, compared to its original version (Guerreri et al., 2021).

Diffusion tensor imaging (DTI) is another widespread MRI technique that has been used to assess neural plasticity in the past (Basser & Pierpaoli, 1996). Unlike NODDI, DTI considers each voxel as single compartment, giving only a composite picture of the tissues. The compartment is modelled by a tensor that describes the 3D profile of diffusion. DTI derived metrics are the mean diffusivity (MD), which quantifies the average diffusivity (the “size” of the tensor) in each voxel, and the fractional anisotropy (FA), quantifying the overall amount of directional preference of diffusion in each voxel (the “shape” of the tensor).

##### 2.3.2.3 Model fitting

The NODDI model was fitted to all available diffusion data (b = 0, 650, 1000, 2000 s/mm2) via a modified version of the NODDI-toolbox (http://mig.cs.ucl.ac.uk/index.php?n=Tutorial.NODDImatlab) implemented in Matlab (The Math Works, Inc. MATLAB. Version 2019b). For a robust parameter estimation (Jelescu et al., 2016; Zhang et al., 2012), the intra-neurite bulk diffusivity was fixed to the value of 2.28 µm2/ms, which is supported by previous studies using a non-conventional, generalized version of dMRI acquisitions (Guerreri et al., 2020). DTI was fitted to the data with b-values up to 1000 s/mm2 via FSL’s DTIFIT tool.

##### 2.3.2.4 Intra-subject group alignment and diffusion to anatomical space registration

To define intra-subject correspondence across DW images from different timepoints, we implemented a robust approach to minimize longitudinal image processing biases such as interpolation asymmetry (Yushkevich et al., 2010). In brief, the approach finds a subject-specific target space that represents the centre of all the timepoints. This approach has the advantage of treating all timepoints equally, which is an essential requirement in longitudinal image processing (Reuter et al., 2012).

Next, for each subject, we defined the correspondence transformation from each of the DWI timepoint and the corresponding T1w image available. This is done in three two steps: first, we determined a subject-specific DW template into the timepoint target space defined previously, which was representative of all timepoints; second, we registered the template to the T1w image via an affine transformation; third, we found the transformation that maps each DWI into the corresponding anatomical image by composing the two transformations.

Finally, all the dMRI derived metrics were resampled from each individual timepoint space into the corresponding anatomical space in one single step to minimize interpolation effects.

##### 2.3.2.5 Inter-subject group correspondence

FreeSurfer software was used to establish the correspondence between subjects by segmenting sub-cortical areas (Fischl et al., 2002, 2004) as well as registering each subject cortical surface to a template surface (Dale et al., 1999; Fischl & Dale, 2000). The pre-processed anatomical images underwent FreeSurfer routine. A visual quality assessment was performed at the end of each of the three main sections of the routine. Ad hoc interventions were carried out when necessary to ensure good quality of the segmentations and of the surface placement. The hippocampus is known to have an important role in spatial navigation (Epstein et al., 2017). In particular, different sub-structures are believed to have different functions (Harland et al., 2017). To account for this, we used FreeSurfer segmentHA_T1 routine (Iglesias et al., 2015) to further divide the segmented hippocampi into head, body and tail.

We assessed sub-cortical GM plasticity-dependent microstructural changes by comparing the dMRI-derived metric average values in each of the considered ROIs across timepoints and subjects. For each subject, the mean values were computed into each timepoint native diffusion space to minimize any possible resampling bias. The FreeSurfer derived segmentations were resampled into each timepoint space inverting the transformation used to bring the dMRI derived maps into the anatomical space. Based on prior navigation studies reviewed above, our subcortical ROI analysis focused on five bilateral regions, namely left and right head, body, and tail of the hippocampus, putamen and caudate.

In the cortical GM, we assessed plasticity-dependent microstructural changes vertex-wise via a surface-based analysis. To increase the sensitivity to subtle changes, it is common practice to perform spatial smoothing which increases the statistical power (Brodt et al., 2018; Tavor et al., 2020). In a surface-based analysis framework the smoothing step occurs on the surface. Hence, compared to conventional voxel-wise volumetric approaches, this procedure has two main advantages: first, it preserves cortical folding spatial continuity i.e., it prevents mixing of information from cortical regions which are spatially close but topologically distant; second, it prevents mixing of information from non-cortical regions such as WM and CSF. In order to implement the surface-based analysis we projected each of the dMRI derived metrics which were resampled into the anatomical space onto the cortical surface. Specifically, to reduce partial volume contamination (Fukutomi et al., 2018), the mid-thickness surface was used. The mid-thickness surface was obtained by expanding the pial surface i.e., the surface marking the boundary between white matter (WM) and grey matter (GM), midway through the cortical thickness via FreeSurfer’s “mris_expand” routine. Finally, the projected maps were resampled into the template surface space. Smoothing with a 6mm FWHM Gaussian filter was performed to all the surface maps in this space before statistical analysis.

#### 2.3.3. Statistical analyses (dMRI data)

All analyses were conducted in the general linear model (GLM) framework for each of the three diffusion imaging parameters of interest (i.e., mean diffusivity [MD], neurite density index [NDI], free water fraction [FWF]). First, we investigated learning-related short-term changes from before (DWI-1) to after (DWI-2) the Day 1 Learning session in the whole sample (N = 32), as well as potential correlations between the Navigation Performance scores and learning-related changes. Second, we investigated potential post-training sleep effects running analysis with between-subject factor Group (RS vs. SD) and within-subject factor either Pre-Learning sessions, i.e., first DWI scan at Day 1 (DWI-1) and Day 4 (DWI-3), or Relearning-related Session (DWI-3 vs. DWI-4). Between-group differences in correlations between learning-related changes and Navigation Performance Scores were also tested.

In the cortical ribbon, contrasts were computed using FreeSurfer’s (http://surfer.nmr.mgh.harvard.edu/) surface-based stream applying a vertex-based approach. Cluster-wise correction for multiple comparisons (p<.05) was performed running a non-parametric permutation simulation using FreeSurfer’s *mri_glmft-sim* command with 10,000 simulations and a vertex wise cluster-forming threshold of p<.001. Peak coordinates of significant clusters are reported based on the MNI system. For the sake of completeness, clusters associated with learning at the uncorrected level (p<.001) are reported as Supplementary Material.

Based on the FreeSurfer volume-based subcortical segmentation stream, we also examined the abovementioned contrasts on MD, NDI and FWF values in our five subcortical left and right regions-of-interest (ROIs) identified as key actors of the navigation learning network (Epstein et al., 2017): the head, body and tail of the hippocampus, the caudate nucleus and the putamen. Results on a priori ROIs are reported at p-level < .05, uncorrected.

## 3. Results

### 3.1. Behavioural Results

#### 3.1.1. Sleep Data

Sleep duration and quality were estimated by means of self-reports of the nights preceding Day 1 and Day 4. Mean sleep duration was not significantly different between RS and SD groups on the night before Day 1 (RS = 439 ± 63 min, SD = 453 ± 41 min; p > 0.2) and the night before Day 4 (RS = 437 ± 55 min, SD = 458 ± 44 min; p > 0.2). Within each group, duration of each night did not differ significantly (RS, p > 0.8; SD, p > 0.7). Likewise, a similar analysis conducted on subjective sleep quality assessed by means of a 6-point scale failed to evidence between-group differences. Subjective sleep quality was equivalent between groups for each night (night before Day 1: RS = 3.7 ± 1.2, SD = 3.5±1.1; p>0.5; night before Day4: RS = 3.6±0.7, SD = 3.7±0.9; p>0.9). Finally, within-group comparisons did not reveal any significant difference in sleep quality between both nights (RS p > 0.8; SD p > 0.1). These results indicate that sleep patterns observed the night before the SD episode were restored before being scanned again in the delayed session.

#### 3.1.2. Navigation Performance

As a reminder, for each of the 10 place finding tests administered at Day 1 (Immediate retrieval) and Day 4 (Delayed retrieval), participants were instructed to reach, as fast as possible, a given Level 1 target from one designated starting point within 28 s. Navigation performance was estimated as the distance remaining to reach destination at the end of the trial. A two-way ANOVA computed on within-session individual mean scores with group (SD vs. RS) and retrieval session (Immediate vs. Delayed) factors did not reveal any significant main or interaction effect (all ps > 0.1; mean distance score at Day 1 SD = 17.08 arbitrary units [standard deviation 8.23] vs RS = 15.19 [13.32]; Day 4 SD = 17.59 [11.2] vs. RS = 15.32 [12.1]). Likewise, at testing after the second learning session on the extended version of the town (distance to destination after the 60 seconds deadline), mean performance did not statistically differ (t = −0.2; p = 0.8) between RS = 15.7 (19.20) and SD = 17.2 (19.6) conditions.

### 3.3. Diffusion Weighted Imaging (DWI) results

#### 3.3.1. Learning-related structural changes (Day 1; DWI-1 vs. DWI-2)

##### 3.3.1.1. Cortical Ribbon

Learning on Day 1 was associated with significantly decreased MD and FWF in the cortical ribbon (Figure 2 [top panel], Supplementary Table 1.a & 1.b). FWF significantly decreased bilaterally in the occipital pole, middle occipital gyrus, angular gyrus, central sulcus expanding in the precentral gyrus in the left hemisphere, and in the superior part of the precentral sulcus. In the right hemisphere, FWF decreased in the parieto-occipital sulcus expanding in the cuneus and precuneus, the posterior transverse collateral sulcus, the lateral occipitotemporal sulcus, in three clusters in the superior temporal sulcus, in the posterior dorsal cingulate gyrus and in the middle frontal gyrus. In the left hemisphere, clusters were located in the middle occipital sulcus, occipito-superior and transversal sulci, in the lingual and fusiform gyri, the intraparietal sulcus, the precuneus, the posterior part of the lateral fissure, the superior part of the precentral and the pars marginalis of the cingulate sulcus. MD similarly decreased in the same locations than FWF, to the exception of one cluster in the left superior temporal sulcus. No significant changes were found for NDI.

**Figure 2.**
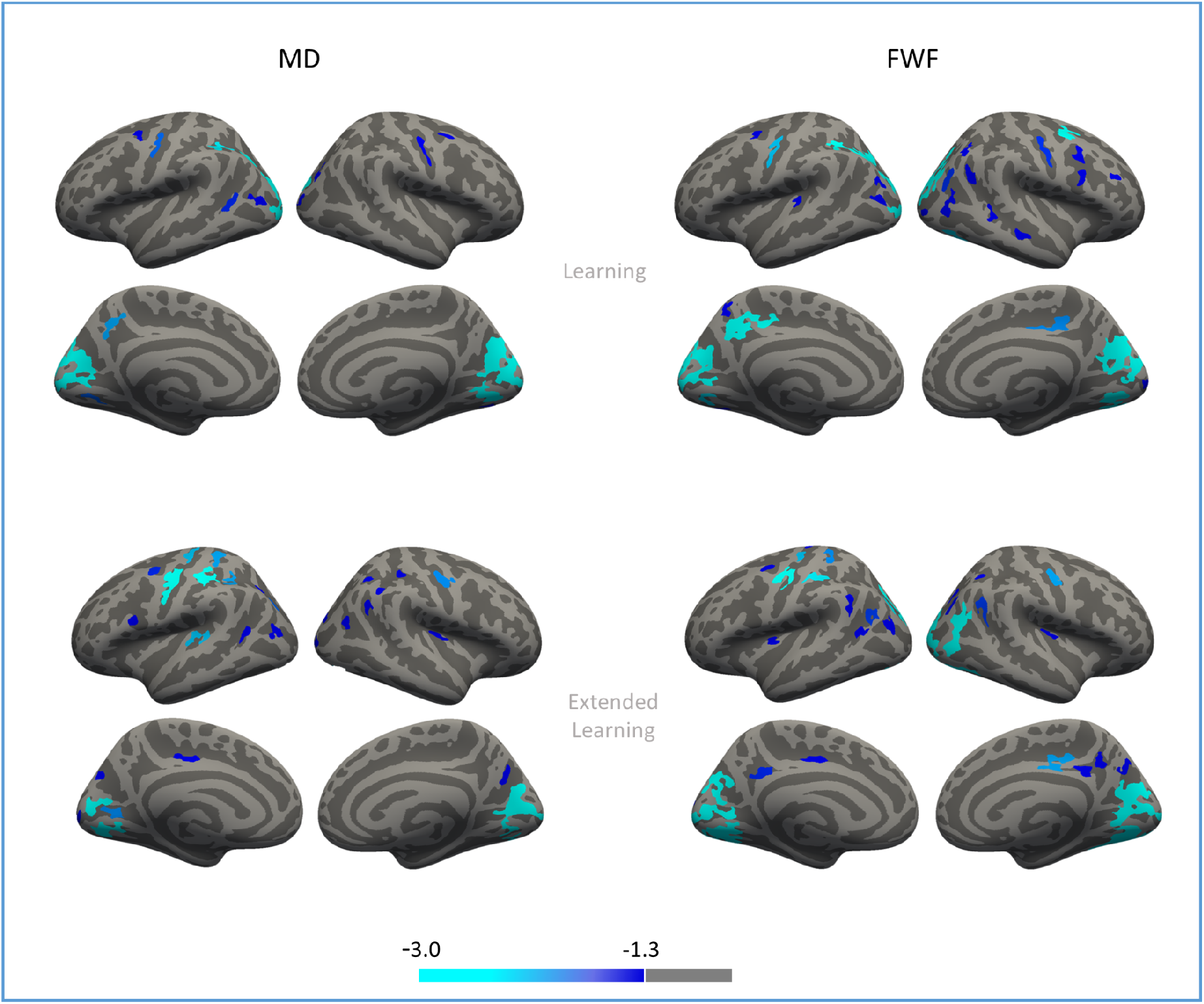
Learning- and Extended Learning-related structural changes (cortical ribbon). Significant clusters of decreases in MD and FWF after Learning on Day 1 (upper panel) and Extended Learning on Day 4 (lower panel) in the cortical ribbon after cluster-wise correction for multiple comparisons. Significance levels are on a -log*(p)* scale. See Supplemental Figure S1 for results without cluster-wise correction for multiple comparison on MD, FWF and NDI parameters.

Day 1 navigation performance scores correlated with MD changes in a post-central gyrus cluster (MNI: x = 43 ; y = -19 ; z = 39; Figure 3). The lower the navigation performance at Day 1, the more MD decreased from the pre- to the post-learning session. No correlation was found with FWF.

**Figure 3.**
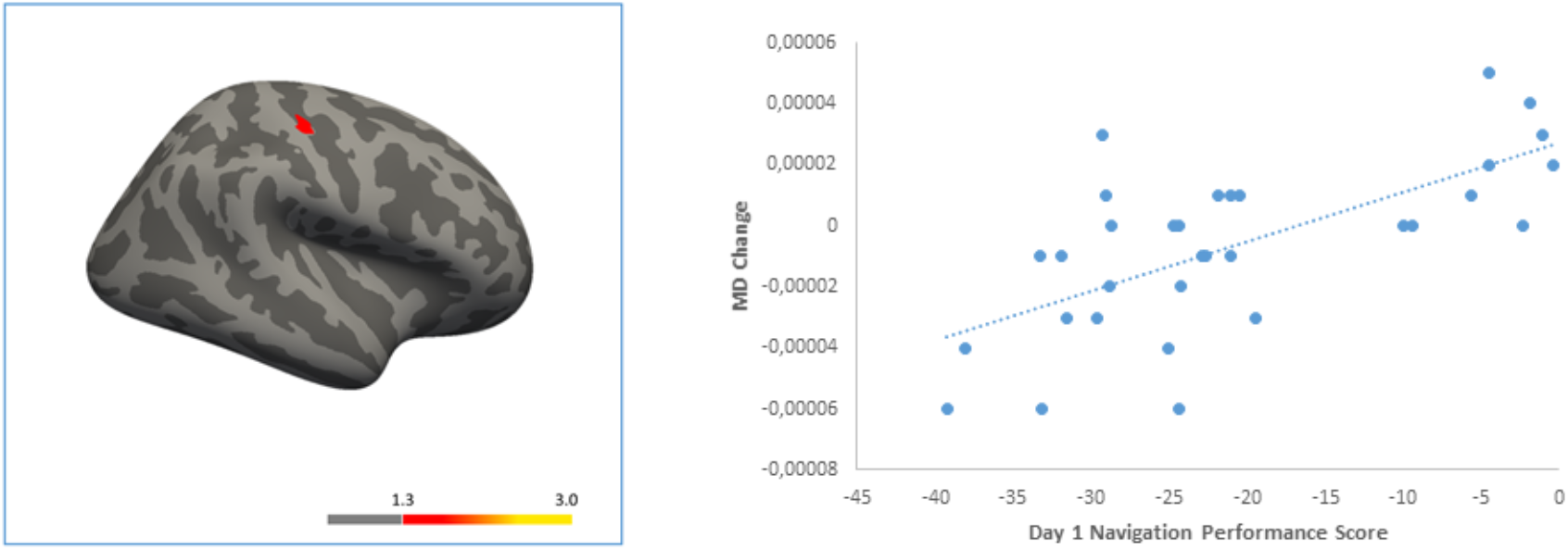
Correlation between performance and structural changes. Left: post-central gyrus cluster (MNI x y z coordinates 43 -19 39 mm) with a positive correlation between MD change and Day 1 navigation performance, after cluster-wise correction for multiple comparisons. Significance level is on a -log*(p)* scale. Right: scatterplot showing the relationship between Day 1 navigation performance scores (computed as the mean distance remaining toward targets in space units, such as higher scores correspond to higher performance levels) and MD change from pre- to post-learning (post-minus pre-learning values, such as positive values correspond to increases in MD over time).

##### 3.3.1.2. Subcortical ROIs

In subcortical ROIs, significant changes were found in the right Caudate, in which FWF decreased by 1.9% (DWI-1 mean(SD) = .1284(.0243) vs. DWI-2 mean(SD) = .1260(.0238), t = 2.09, p = .044, g = .1) and NDI decreased by 0.8% (DWI-1 = .3453(.0329) vs. DWI-2 = .3426(.0321), t = 2.3, p = .029, g = .08), and in the left and right Body of the Hippocampus, in which FWF decreased by 1.9% and 1.7% (Left: DWI-1 = .1672(.0282) vs. DWI-2 = .1641(.0281), t = 2.38, p = .023, g = .11 ; Right: DWI-1 = .1839(.0220) vs. DWI-2 = .1807(.0243), t = 2.08, p = .045, g = .13).

Day 1 navigation performance scores correlated with MD changes in the right caudate (r = .42, p = 0.016) and putamen (r = -.39, p = .027), with FWF changes in the left putamen (r = -.42, p = .16), and with NDI changes in the left putamen (r = .48, p = .02).

#### 3.3.2. Sleep-related effects in Pre-learning vs. Pre-relearning (DWI-1 vs. DWI-3 X RS vs. SD)

No significant effect was found in the cortical ribbon for any of the three indices. A main effect of sleep was detected in the subcortical ROIs, where a significant change was found in the right Putamen; NDI decreased by 0.8% and 0.7% (Noddi1, mean(SD): .3942(.0270), Noddi3, mean(SD): .3915(.0285), t = 2.63, p = .013, g = .09). No significant correlations between mean changes and Day 1 performance scores were found. Significant correlations between mean changes and Day 4 performance scores were found in the right Caudate for MD (r = .36, p = .047), in the left and right Putamen for FWF (Left: r = - .36, p = .047; Right: r = -.46, p = .01).

No significant interaction effects were found either in the cortical ribbon or in the subcortical ROIs, for any of the three indices. No difference in correlation with Day 4 navigation performance was found between the two groups.

#### 3.3.3. Sleep-related effects on Extended Learning (DWI-3 vs. DWI-4 X RS vs. SD)

##### 3.3.3.1. Cortical Ribbon

The analysis evidenced a main session effect (DWI-3 vs DWI-4), but no Group (RS vs. SD) nor Group X Session interaction effects in the cortical ribbon. Regarding the main session effect, extended learning on Day 4 was associated with significantly decreased MD and FWF in the cortical ribbon (Figure 2 [bottom panel], Supplementary Table 2.a & 2.b). FWF decreased bilaterally in the occipito-superior and transversal sulcus, the parieto-occipital sulcus, the superior temporal and the central sulcus. In the right hemisphere, clusters were localized in the lingual gyrus, the cuneus gyrus, the middle occipital gyrus, the superior occipital gyrus, the subparietal sulcus, the supramarginal gyrus, the posterior cingulate sulcus and gyrus, the postcentral gyrus and sulcus, the precentral sulcus and the circular sulcus. In the left hemisphere, clusters were found in the occipital pole, the calcarine sulcus, the middle occipital sulcus and gyrus, the superior occipital gyrus, the subparietal sulcus, the intraparietal and transversal parietal sulcus, the posterior part of the lateral fissure and in the pars marginalis of the cingulate sulcus. Clusters found significant for FWF encompassed all clusters significant for MD to the exception of one cluster in the central sulcus (x = -14.5 ; y = -29.1 ; z = 58.9).

No significant correlations with Day 4 navigation performance scores were found for any of the three indices, as well as no difference in correlation between RS and SD conditions.

##### 3.3.3.2. Subcortical ROIs

The analysis evidenced main Session (DWI-3 vs DWI-4) and Group by Session interaction effects, but no main Group (RS vs. SD) effect. Group by Session interaction effects on FWF were found in the right Caudate (RS 2.7% decrease = -.0033 [.0050] vs. SD 0.9% increase = +.0011 [.0062]; t = 2.26, p = .031), and on NDI in the right Caudate (RS 0.2% decrease = -.0007[.0057] vs. SD 1.3% increase = +.0045 [.0062]; t = 2.46, p = .02) and the left Head of the Hippocampus (RS 1.3% decrease= -.0029 [.0076] vs. SD 1.8% increase = .0039 [.0087]; t = 2.35, p = .025) (Figure 4).

**Figure 4.**
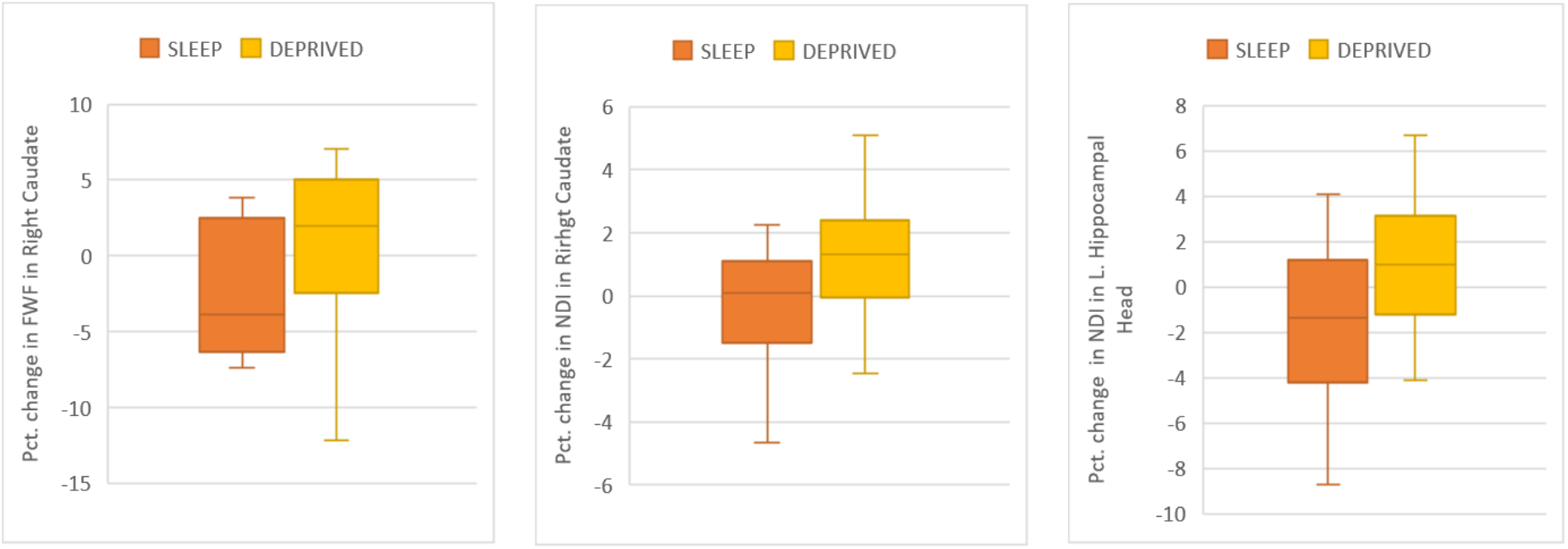
Sleep-related effects on Extended Learning (DWI-3 vs DWI-4 vs RS vs SD). Box Plot showing sleep-related changes in the Sleep vs. Sleep Deprived group in percentage (with standard deviation) for a) Free Water Fraction (FWF) in the Right Caudate (RS 2.7% decrease vs. SD 0.9% increase), b) Neurite Density Index (NDI) in the Right Caudate (RS 0.2% decrease vs. SD 1.3% increase) and c) NDI in the Left Hippocampal Head (RS 1.3% decrease vs. SD 1.8% increase).

Regarding the main Session effect (i.e., relearning-related changes), MD decreased over time by 0.4% in the right Caudate (DWI-3 mean(SD) = .0007296(.00001351) vs. DWI-4 mean(SD) = .0007265(.00001221), t = 2.78, p = .009, g = .23), by 0.5% in the Left and Right Body of the Hippocampus (Left Hpc DWI-3 = .0007882(.00001780) vs. DWI-4 = .0007843(.00001713), t = 2.09, p = .044, g = .22; Right Hpc DWI-3 = .0008015(.00001913) vs. DWI-4 = .0007975(.00001805), t = 2.14, p = .04, g = .21) and by 0.4% in the right Hippocampus Head (DWI-3 = .0008048(.00001206) vs. DWI-4 = .0008012(.00001094), t = 2.67, p = .012, g = 0.3). FWF decreased by 1.5% in the left Hippocampus Head (DWI-3 = .1647(.0243) vs. DWI-4 = .1622(.0230), t = 2.19, p = .036, g = .1), by 2.3% and 2.5% in the Left and Right Body of the Hippocampus, respectively (Left Body DWI-3 = .1685(.0298) vs. DWI-4 = .1647(.0298), t = 2.85, p = .008, g = .12; Right Body DWI-3 = .1851(.0243) vs. DWI-4 = .1805(.0254), t = 3.31, p = .002, g = .18), and by 2.6% and 3.1% in the Left Right Tail and of the Hippocampus (Left Tail DWI-3 = .1755(.0411) vs. DWI-4 = .1709(.0405), t = 2.77, p = .009, g = .11; Right Tail DWI-3 = .1835(.0413) vs. DWI-4 = .1779(.0427), t = 3.99, p < .001, g = .13). Finally, NDI increased by 0.6% in the Left Putamen (DWI-3 = .3954(.0246) vs. DWI-4 = .3979(.0245), t = - 2.19, p = .036, g = -.1).

Significant correlations between pre- to post-relearning NDI changes and Day 4 performance scores were found in the left Head of the Hippocampus (r = .43, p = .017) and in the left Putamen (r = .47, p = .009). No difference in correlation with Day 4 navigation performance was found between the two groups for any of the three indices.

## Discussion

In the present study, we used advanced dMRI analysis techniques to investigate the offline microstructural changes associated with learning and extended learning following sleep in a spatial navigation task.

In line with previous studies (e.g., Orban et al., 2006; Rauchs, Orban, Schmidt, et al., 2008), behavioural performance was similar between the sleep deprived (SD) and regular sleep (RS) group, both after initial learning, and when tested after exploring an extended version of the town. Our behavioural results suggest that (1) both groups gained knowledge of the virtual town that persisted 3 days after learning, that (2) SD on the first post-training night did not overtly alter subjects’ ability to find their way in the town at delayed retrieval (which does not preclude the use of qualitatively different neural and cognitive strategies for a similar performance), and that (3) learning novel routes and integration with prior knowledge in the extended version of the town was quantitatively similar in the RS and SD conditions.

In the whole sample, learning and extended learning episodes were associated with an overall pattern of MD (DTI) and FWF (NODDI) decreases in the cortical ribbon. Maps of MD and FWF changes were highly coherent (see Supplementary figure 1). As a reminder, the DTI mean diffusivity (MD) parameter quantifies the average diffusivity in each voxel. MD was proposed in previous studies to be a relevant and sensitive marker of tissue microstructure changes associated with learning. However, the biological interpretation of MD changes is not straightforward. Indeed, the calculated tensor is an average of all cellular components within the measured voxel, which can be affected by several parameters of tissue density or tissue organization, such as synaptogenesis, changes in the morphometry of axons, dendrites and glial processes, or alterations in cell body size and shape (Sagi et al., 2012). In the NODDI model, FWF is the component that accounts for the presence of cerebrospinal fluid or oedema in the tissues volume, besides the intra- and extra-neurite spaces (Zhang et al., 2012). Reduced FWF thus theoretically entails more space occupied by the two other compartments. Since the intra-neurite compartment signal fraction NDI parameter was unchanged at the cortical level, our results suggests that structural modifications took place in the extra-neurite space including glial cells, in line with the interpretation proposed in Tavor et al. (2013) based on the CHARMED framework. Astrocytes are key elements of synaptic function, and exhibit rapid changes following synaptic activity (Theodosis et al., 2008). Evidence of structural plasticity in astrocyte morphology was previously found in rats trained on the Morris water maze (Blumenfeld-Katzir et al., 2011). Rapid structural modifications are also possible in dendritic spines, as formation of dendritic spines after training was shown to occur within the hour (Xu et al., 2009).

Both navigation learning and relearning episodes evoked overall similar patterns of microstructural changes, that are in line with fMRI findings in spatial navigation studies (Epstein et al. 2017). On the medial surface, clusters were mostly localised in the occipital and parietal lobes, while clusters found on the lateral surface were mostly localised in the occipital, parietal, temporal and posterior frontal areas (see Supplementary figure 1). On day 1, navigational performance positively correlated with MD change in the post-central gyrus, that comprises the primary somatosensory cortex. While spatial representation in the brain has been largely associated with the hippocampus and the entorhinal cortex, one recent study identified place cells, head direction cells, boundary vector/border cells, grid cells and conjunctive cells, in the primary somatosensory cortex of rats (Long & Zhang, 2021). These newly identified somatosensory spatial cells form a spatial map outside the hippocampal formation, support the hypothesis that location information modulates body representation in the somatosensory cortex, and that spatial tuning in the primary somatosensory cortex may be crucial for body representation/simulation in space. Here, MD reductions in the post-central gyrus were more pronounced in participants with lower performance, suggesting that increased engagement of this brain area was associated with less efficient learning.

In subcortical ROIs, learning and relearning were associated with a set of changes in MD and FWF again, but also in NDI. In the hippocampus, MD decreases were found for the extended learning episode only, whereas FWF decreased both in learning and extended (re)learning episodes. These findings concur with previous studies that associated spatial navigation learning with MD decreases in hippocampal areas (Keller & Just, 2016; Sagi et al., 2012; Tavor et al., 2013). A discrepant finding is the absence of decreased MD in the first learning episode. Besides possible statistical power effects, it should be acknowledged that the spatial navigation task we used was different from previous studies using dMRI to explore the microstructural changes associated with spatial learning. Indeed, it is likely more complex as participants have to learn to find their way in a maze, as opposed to learning to repeatedly drive a single road. As for other subcortical ROIs, FWF reductions were found in the right caudate after the learning episode, and MD reductions after the extended learning episode. Finally, NDI decreased in the right caudate after learning, but increased in the left putamen after relearning. Also, positive correlations were found between the NDI parameter and navigational performance in the left putamen both at learning and relearning. Increased NDI at relearning and positive correlation with performance may reflect a plasticity-related restructuration of the intra-neurite space bounded by neurite membranes and myelin sheaths, consistent with concomitantly reduced FWF.

Regarding sleep-related consolidation effects, no differential changes were found in the cortical ribbon at extended learning when comparing the RS and SD conditions. Besides statistical power issues, this null finding may indicate that sleep-dependent structural reorganization at the cortical level was not yet implemented and/or detectable up to 3 days after the sleep manipulation, in line with fMRI findings that hippocampo-neocortical transfer of information was not observed two days after sleep deprivation but only weeks to month later in young adults (Gais et al., 2007). At variance, in subcortical ROIs, microstructural changes associated with extended learning were modulated by post-learning sleep. In the left hippocampus head, NDI decreased in participants allowed to sleep after learning (RS) but increased in the sleep-deprived ones (SD). Also in the right caudate nucleus, NDI increased in the SD condition. Finally, FWF decreased in the right Caudate in the RS condition and increased in the SD condition. Hence, our data suggest that sleep following learning influences the microstructural underpinning of later extended learning episodes in which novel information is integrated into existing knowledge, in such a way that opposite patterns can be detected in key areas supporting spatial navigation learning. Decreased hippocampal NDI paralleled by stabilized NDI (and reduced FWF) in the caudate in the RS condition may to some extent reflect the gradual shift from a hippocampus-dependent spatial strategy to a more integrated and automated striatal response-based strategy, as found in fMRI studies (Orban et al., 2006; Rauchs, Orban, Schmidt, et al., 2008).

One potential limitation of the present study is the lack of a control condition investigating the microstructural changes associated with exposure to spatial navigation without learning. Including such a control condition may have helped identifying clusters of changes playing a key role in learning and extended learning, as opposed to brain areas involved in spatial navigation. Another potential limitation is the fact that our ROI analysis did not correct for the examination of multiple regions of interest, increasing the risks of type 1 error despite the fact that we worked on a priori defined ROIs. For these reasons, we consider our ROI-based results as preliminary. On the other hand, the investigation, of learning and extended learning changes in the cortical ribbon was corrected for multiple comparisons. Also on the positive side, our processing pipeline was optimized separately for cortical and sub-cortical GM analysis. Previous studies (e.g., Brodt et al., 2018; Sagi et al., 2012; Tavor et al., 2013, 2020) used voxel-based non-linear registration to define correspondence across subjects, combined with spatial smoothing to increase statistical power. We believe that such an approach may not be optimal for at least two reasons. First, volumetric smoothing entails the risk of mixing information from non-GM regions such as CSF and WM/ Scond, in the highly convoluted cortical foldings, volumetric smoothing can induce mixing of information from cortical regions which are spatially close but topologically distant. In the present study, we used two specific analysis streams for cortical and subcortical GM analyses, aimed at avoiding such limitations. In the surface-based analysis in particular, the smoothing step occurs on the surface after the projection of the relevant information from the cortical GM voxels is performed, hence preventing information mixing as described above. For its part, ROI analysis does not involve any smoothing step and thus should not be affected by the reported information mixing issue.

To sum up, our results disclose navigation learning-related decreased dMRI parameters in the cortical ribbon and subcortical areas. Post-learning sleep-related changes were found in the extended learning session in the caudate nucleus and the hippocampus, suggesting a sleep-related remodelling of neurites and glial cells subtending learning and memory processes in basal ganglia and hippocampal structures, that may precede further reorganisation at the cortical level.

## Acknowledgments

This study was supported by the Fonds de la Recherche Scientifique (F.R.S.-F.N.R.S.) and the Fonds Wetenschappelijk Onderzoek – Vlaanderen (F.W.O.) under the Excellence of Science (EOS) Project (MEMODYN, No. 30446199). W.S. is supported by F.N.R.S. Aspirant Research Fellowship. M.G. received postdoctoral salary support from the EOS grant. At the time of the study, T.V. was funded by an Université Libre de Bruxelles (ULB) Individual Fellowship. M.D. is supported by the Fonds pour la Recherche dans l’Industrie et l’Agriculture (FRIA) Fellowship. C.S. is FNRS Research Associate.

## SUPPLEMENTARY MATERIAL

**Supplementary Figure 1.**
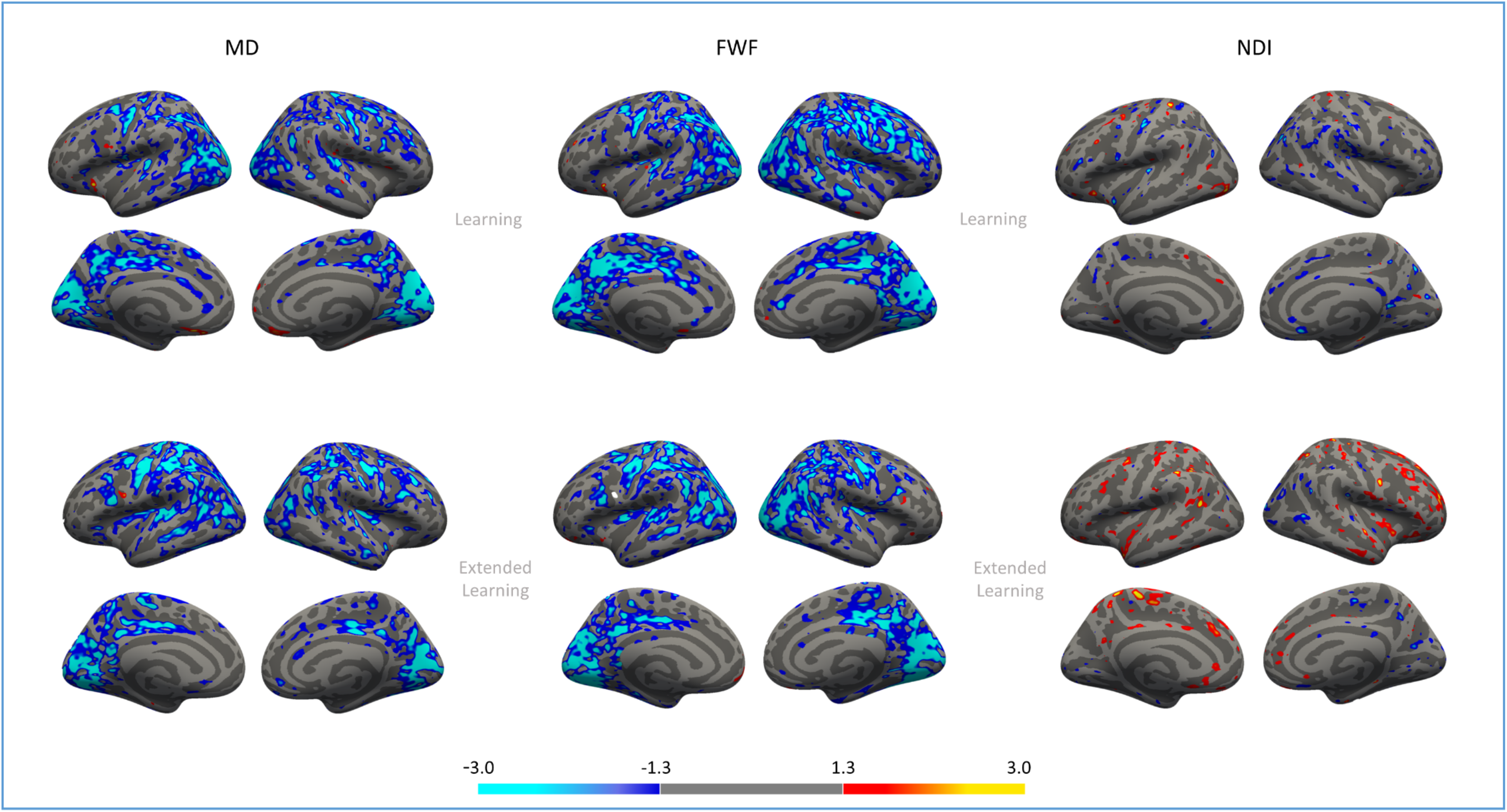
Clusters showing significant changes in MD, FWF and NDI after Learning on Day 1 (upper-part), and Extended Learning (lower-part) on Day 4, uncorrected for multiple comparisons.

**Supplementary Figure 2.**
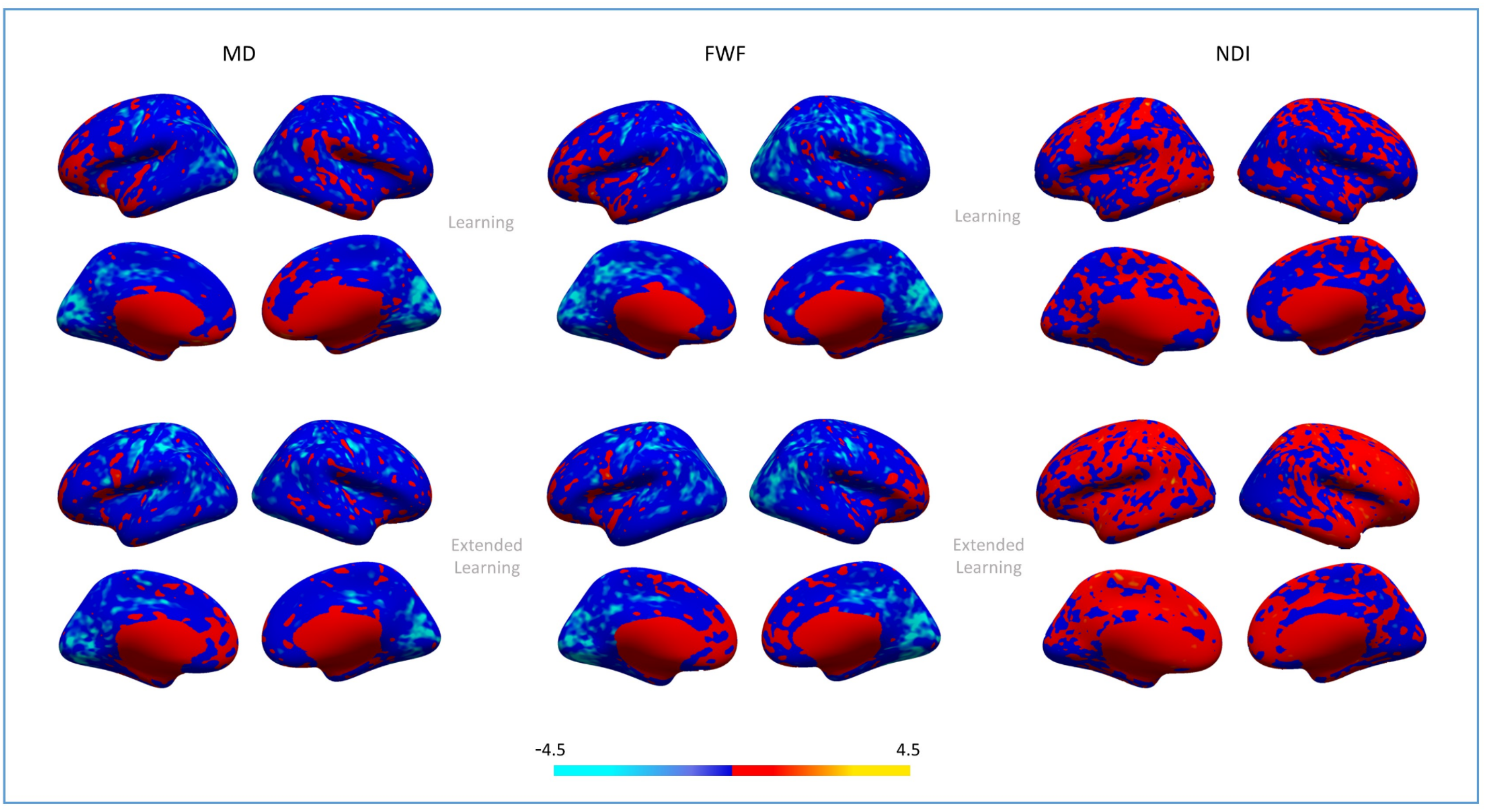
T-statistic maps for the Day 1 (Post- minus pre-learning, upper part) and for the Day 4 (Post-minus –pre relearning, lower part) contrats, for MD, FWF and NDI.

**Supplementary Figure 3.**
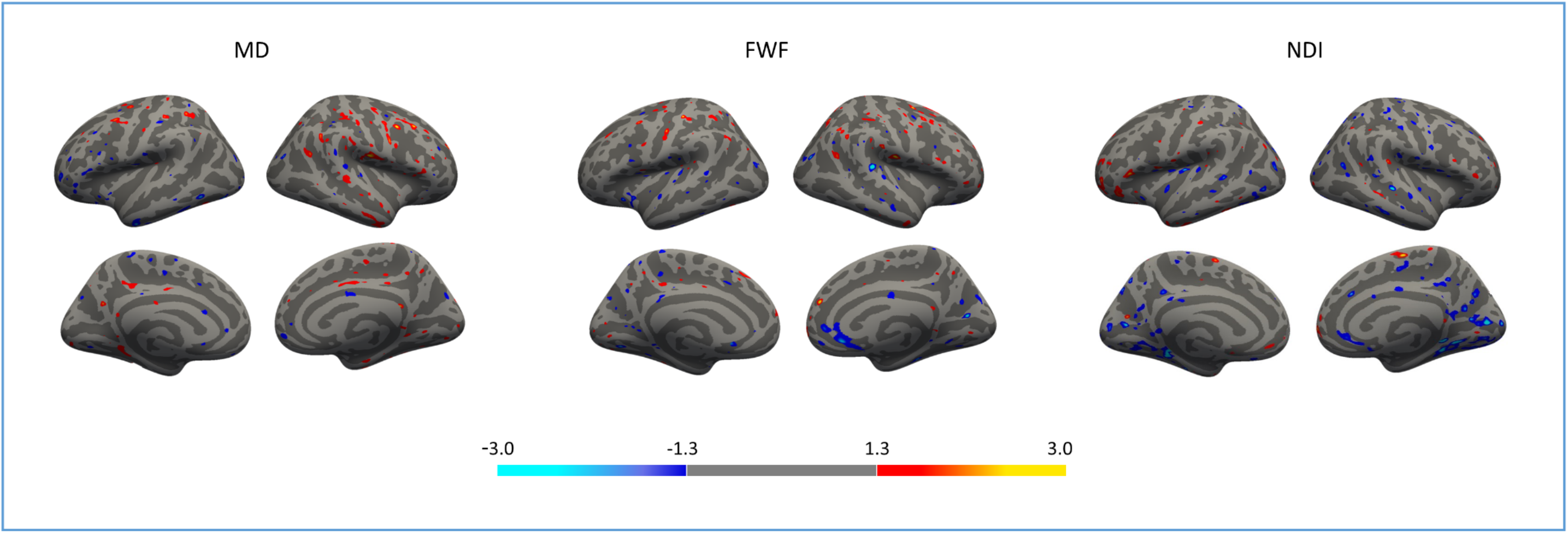
Clusters showing differences in MD, FWF and NDI change between the Sleep and Sleep Deprived groups on Day 4 (Extended Learning), uncorrected for multiple comparisons. Red clusters indicate higher changes for the Post minus Pre Relearning contrast in the Sleep Deprived group when compared with the Sleep group.

**Supplementary Table 1.**
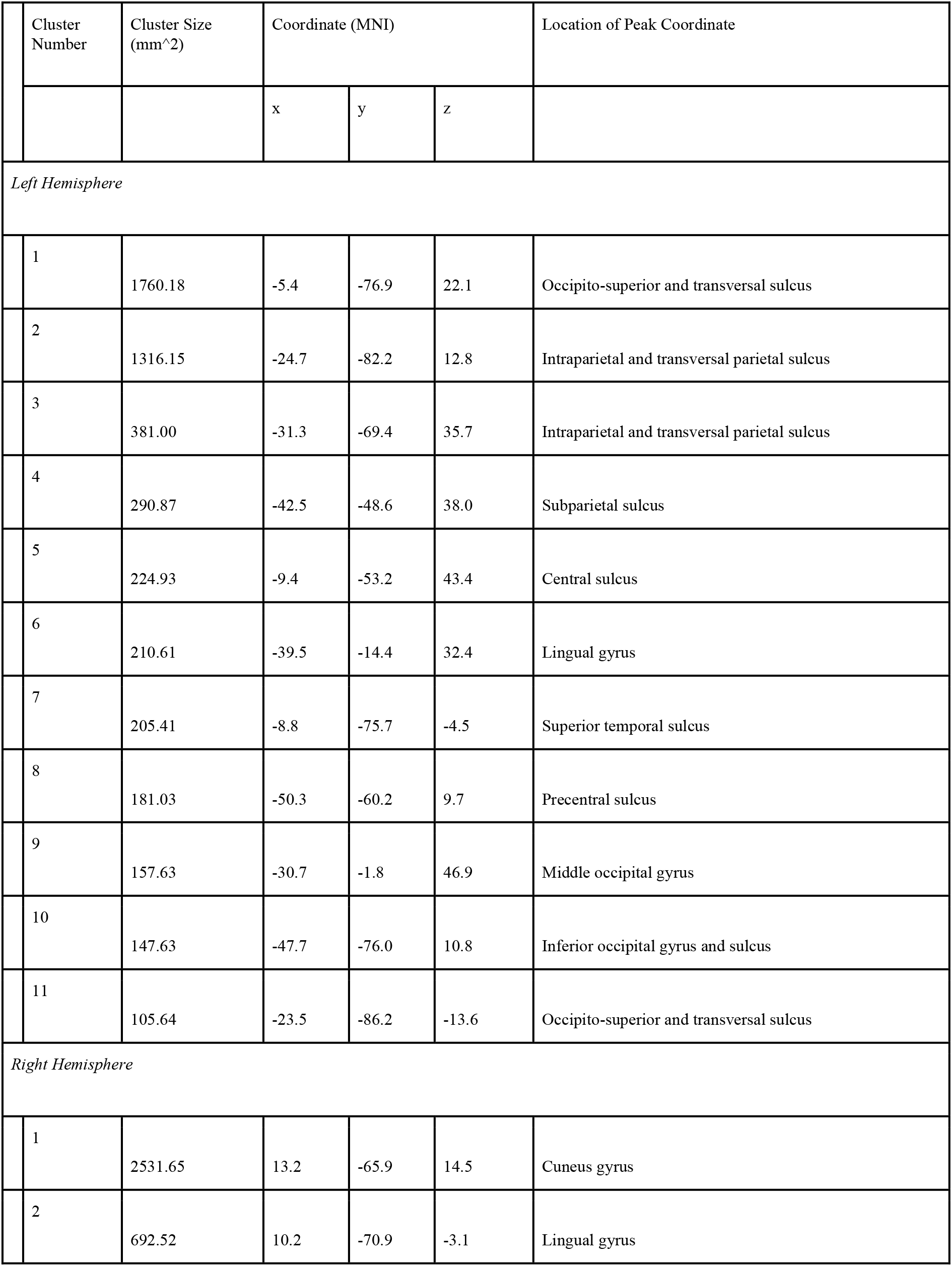
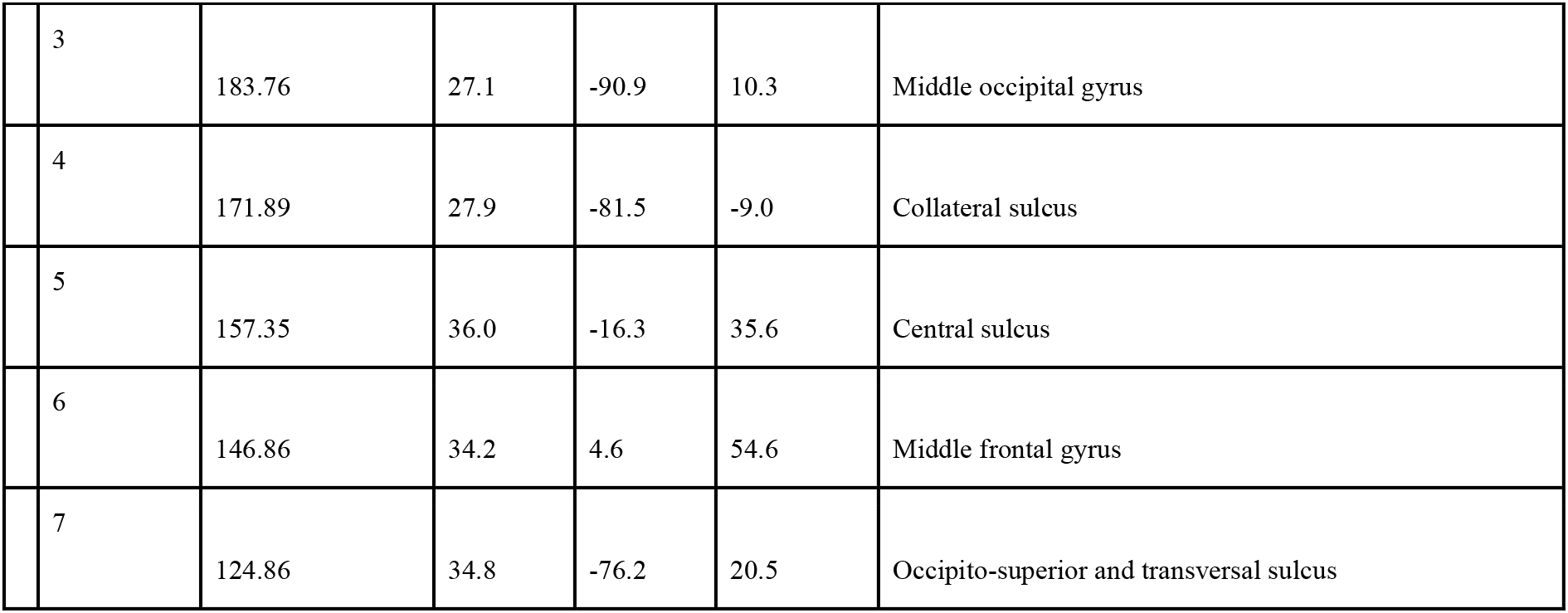
a. List of left and right hemisphere clusters with significant decreases in MD after Learning on Day 1, after cluster-wise correction for multiple comparisons.

|  | Cluster Number | Cluster Size (mm<sup>2</sup>) | Coordinate (MNI) |  |  | Location of Peak Coordinate |
| --- | --- | --- | --- | --- | --- | --- |
|  |  |  | x | y | z |  |
| *Left Hemisphere* |  |  |  |  |  |  |
|  | 1 | 1760.18 | -5.4 | -76.9 | 22.1 | Occipito-superior and transversal sulcus |
|  | 2 | 1316.15 | -24.7 | -82.2 | 12.8 | Intraparietal and transversal parietal sulcus |
|  | 3 | 381.00 | -31.3 | -69.4 | 35.7 | Intraparietal and transversal parietal sulcus |
|  | 4 | 290.87 | -42.5 | -48.6 | 38.0 | Subparietal sulcus |
|  | 5 | 224.93 | -9.4 | -53.2 | 43.4 | Central sulcus |
|  | 6 | 210.61 | -39.5 | -14.4 | 32.4 | Lingual gyrus |
|  | 7 | 205.41 | -8.8 | -75.7 | -4.5 | Superior temporal sulcus |
|  | 8 | 181.03 | -50.3 | -60.2 | 9.7 | Precentral sulcus |
|  | 9 | 157.63 | -30.7 | -1.8 | 46.9 | Middle occipital gyrus |
|  | 10 | 147.63 | -47.7 | -76.0 | 10.8 | Inferior occipital gyrus and sulcus |
|  | 11 | 105.64 | -23.5 | -86.2 | -13.6 | Occipito-superior and transversal sulcus |
| *Right Hemisphere* |  |  |  |  |  |  |
|  | 1 | 2531.65 | 13.2 | -65.9 | 14.5 | Cuneus gyrus |
|  | 2 | 692.52 | 10.2 | -70.9 | -3.1 | Lingual gyrus |

|  |  |  |  |  |  |
| --- | --- | --- | --- | --- | --- |
| 3 | 183.76 | 27.1 | -90.9 | 10.3 | Middle occipital gyrus |
| 4 | 171.89 | 27.9 | -81.5 | -9.0 | Collateral sulcus |
| 5 | 157.35 | 36.0 | -16.3 | 35.6 | Central sulcus |
| 6 | 146.86 | 34.2 | 4.6 | 54.6 | Middle frontal gyrus |
| 7 | 124.86 | 34.8 | -76.2 | 20.5 | Occipito-superior and transversal sulcus |

**Supplementary Table 1.** b. List of left and right hemisphere clusters of significant decreases in FWF after Learning on Day 1, after cluster-wise correction for multiple comparisons.

| Cluster Number | Cluster Size (mm^2) | Coordinate (MNI) |  |  | Location of Peak Coordinate |
| --- | --- | --- | --- | --- | --- |
|  |  | x | y | z |  |
| Left Hemisphere |  |  |  |  |  |
| 1 | 2446.08 | -24.5 | -83.4 | 12.6 | Occipito-superior and transversal sulcus |
| 2 | 1007.28 | -23.8 | -64.7 | 28.2 | Intraparietal sulcus |
| 3 | 680.83 | -14.9 | -35.1 | 38.4 | Pars marginalis of the cingulate sulcus |
| 4 | 490.39 | -31.2 | -84.0 | -0.2 | Middle occipital sulcis |
| 5 | 318.46 | -8.7 | -76.0 | -3.7 | Lingual gyus |
| 6 | 244.64 | -47.5 | -7.9 | 25.0 | Central sulcus |
| 7 | 165.11 | -19.6 | -93.2 | -10.2 | Occipital pole |
| 8 | 143.98 | -11.0 | -56.5 | 55.4 | Precuneus gyrus |
| 9 | 134.68 | -30.2 | -1.6 | 47.0 | Precentral sulcus |
| 10 | 128.18 | -29.7 | -70.3 | -12.5 | Fusiform gyrus |
| 11 | 123.41 | -47.3 | -76.5 | 10.8 | Middle occipital gyrus |
| 12 | 115.26 | -40.1 | -74.2 | 25.1 | Angular gyrus |
| 13 | 100.88 | -38.5 | -32.3 | 9.2 | Posterior part of the lateral fissure |
| Right Hemisphere |  |  |  |  |  |
| 1 | 3366.77 | 16.7 | -60.7 | 27.4 | Parieto-occipital sulcus |
| 2 | 1108.36 | 27.0 | -81.8 | -9.2 | Posterior transverse collateral sulcus |
| 3 | 374.91 | 29.5 | -2.6 | 45.4 | Precentral sulcus |
| 4 | 318.89 | 40.1 | -62.8 | -12.0 | Occipito-temporal sulcus |
| 5 | 252.05 | 6.3 | -36.9 | 42.2 | Posterior dorsal cingulate gyrus |
| 6 | 220.78 | 40.7 | -12.6 | 31.1 | Central sulcus |
| 7 | 209.53 | 43.7 | -68.1 | 28.8 | Angular gyrus |
| 8 | 196.15 | 45.1 | -74.1 | 3.2 | Middle occipital gyrus |
| 9 | 180.89 | 35.3 | 5.4 | 35.4 | Precentral sulcus |
| 10 | 160.97 | 45.3 | -61.0 | 44.4 | Angular gyrus |
| 11 | 160.32 | 17.1 | -94.7 | 1.8 | Occipital pole |
| 12 | 141.82 | 44.2 | -55.1 | 23.5 | Superior temporal sulcus |
| 13 | 120.22 | 46.7 | -43.2 | 4.6 | Superior temporal sulcus |
| 14 | 118.54 | 53.8 | -25.0 | -12.8 | Superior temporal sulcus |
| 15 | 109.47 | 42.8 | 36.8 | 25.9 | Middle frontal gyrus |
| 16 | 108.63 | 43.4 | 8.2 | 19.5 | Precentral sulcus |
| 17 | 104.46 | 34.2 | -87.4 | -1.1 | Middle occipital gyrus |

**Supplementary Table 2.**
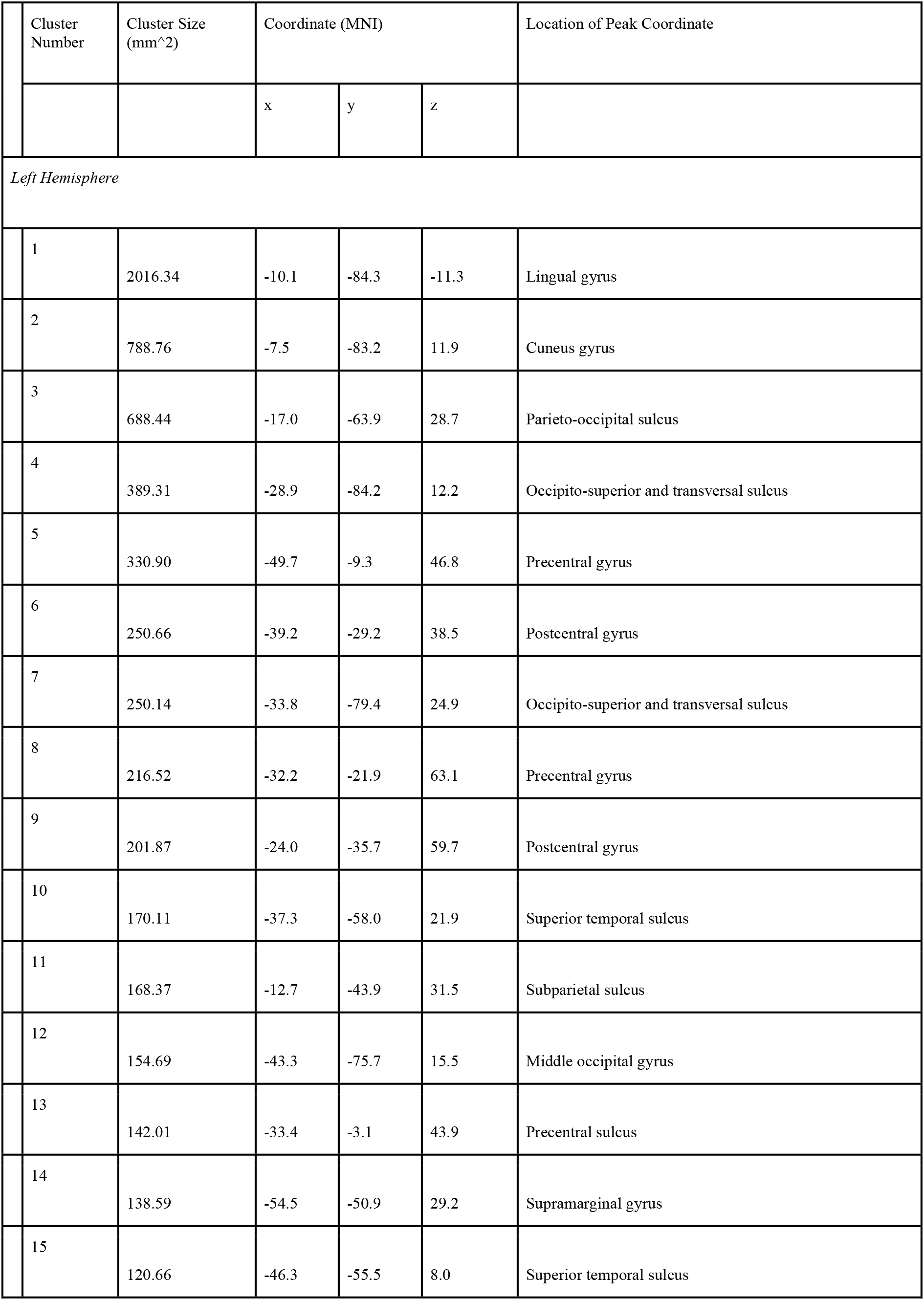
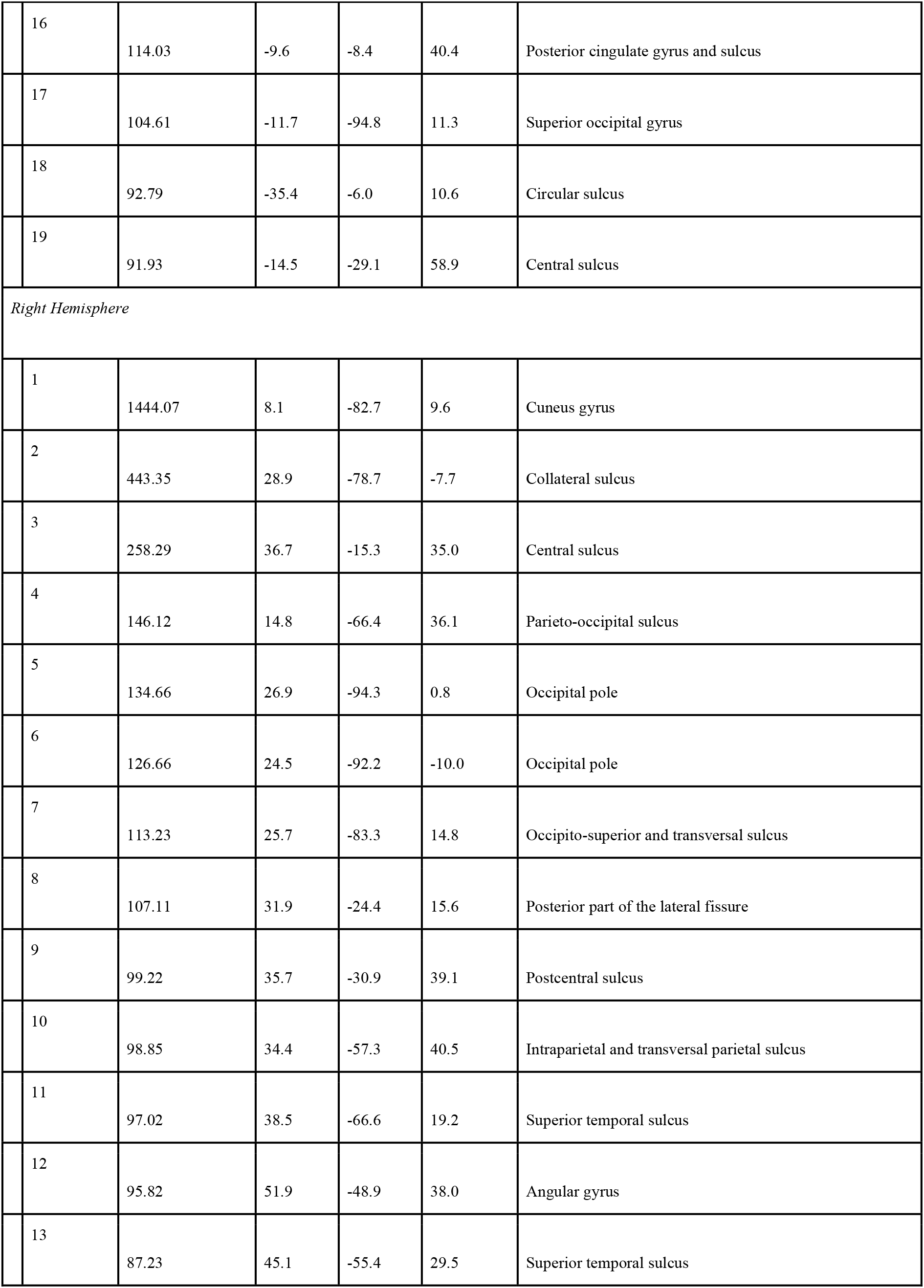
a. List of left and right hemisphere clusters of significant decreases in MD after Learning on Day 4, after cluster-wise correction for multiple comparisons.

|  | Cluster Number | Cluster Size (mm<sup>2</sup>) | Coordinate (MNI) |  |  | Location of Peak Coordinate |
| --- | --- | --- | --- | --- | --- | --- |
|  |  |  | x | y | z |  |
| *Left Hemisphere* |  |  |  |  |  |  |
|  | 1 | 2016.34 | -10.1 | -84.3 | -11.3 | Lingual gyrus |
|  | 2 | 788.76 | -7.5 | -83.2 | 11.9 | Cuneus gyrus |
|  | 3 | 688.44 | -17.0 | -63.9 | 28.7 | Parieto-occipital sulcus |
|  | 4 | 389.31 | -28.9 | -84.2 | 12.2 | Occipito-superior and transversal sulcus |
|  | 5 | 330.90 | -49.7 | -9.3 | 46.8 | Precentral gyrus |
|  | 6 | 250.66 | -39.2 | -29.2 | 38.5 | Postcentral gyrus |
|  | 7 | 250.14 | -33.8 | -79.4 | 24.9 | Occipito-superior and transversal sulcus |
|  | 8 | 216.52 | -32.2 | -21.9 | 63.1 | Precentral gyrus |
|  | 9 | 201.87 | -24.0 | -35.7 | 59.7 | Postcentral gyrus |
|  | 10 | 170.11 | -37.3 | -58.0 | 21.9 | Superior temporal sulcus |
|  | 11 | 168.37 | -12.7 | -43.9 | 31.5 | Subparietal sulcus |
|  | 12 | 154.69 | -43.3 | -75.7 | 15.5 | Middle occipital gyrus |
|  | 13 | 142.01 | -33.4 | -3.1 | 43.9 | Precentral sulcus |
|  | 14 | 138.59 | -54.5 | -50.9 | 29.2 | Supramarginal gyrus |
|  | 15 | 120.66 | -46.3 | -55.5 | 8.0 | Superior temporal sulcus |

|  |  |  |  |  |  |
| --- | --- | --- | --- | --- | --- |
| 16 | 114.03 | -9.6 | -8.4 | 40.4 | Posterior cingulate gyrus and sulcus |
| 17 | 104.61 | -11.7 | -94.8 | 11.3 | Superior occipital gyrus |
| 18 | 92.79 | -35.4 | -6.0 | 10.6 | Circular sulcus |
| 19 | 91.93 | -14.5 | -29.1 | 58.9 | Central sulcus |
| *Right Hemisphere* |  |  |  |  |  |
| 1 | 1444.07 | 8.1 | -82.7 | 9.6 | Cuneus gyrus |
| 2 | 443.35 | 28.9 | -78.7 | -7.7 | Collateral sulcus |
| 3 | 258.29 | 36.7 | -15.3 | 35.0 | Central sulcus |
| 4 | 146.12 | 14.8 | -66.4 | 36.1 | Parieto-occipital sulcus |
| 5 | 134.66 | 26.9 | -94.3 | 0.8 | Occipital pole |
| 6 | 126.66 | 24.5 | -92.2 | -10.0 | Occipital pole |
| 7 | 113.23 | 25.7 | -83.3 | 14.8 | Occipito-superior and transversal sulcus |
| 8 | 107.11 | 31.9 | -24.4 | 15.6 | Posterior part of the lateral fissure |
| 9 | 99.22 | 35.7 | -30.9 | 39.1 | Postcentral sulcus |
| 10 | 98.85 | 34.4 | -57.3 | 40.5 | Intraparietal and transversal parietal sulcus |
| 11 | 97.02 | 38.5 | -66.6 | 19.2 | Superior temporal sulcus |
| 12 | 95.82 | 51.9 | -48.9 | 38.0 | Angular gyrus |
| 13 | 87.23 | 45.1 | -55.4 | 29.5 | Superior temporal sulcus |

**Supplementary Table 2.** b. List of left and right hemisphere clusters of significant decreases in FWF after Extended Learning on Day 4, after cluster-wise correction for multiple comparisons.

|  | Cluster Number | Cluster Size (mm<sup>2</sup>) | Coordinate (MNI) |  |  | Location of Peak Coordinate |
| --- | --- | --- | --- | --- | --- | --- |
|  |  |  | x | y | z |  |
| *Left Hemisphere* |  |  |  |  |  |  |
|  | 1 | 2016.34 | -10.1 | -84.3 | -11.3 | Lingual gyrus |
|  | 2 | 788.76 | -7.5 | -83.2 | 11.9 | Cuneus gyrus |
|  | 3 | 688.44 | -17.0 | -63.9 | 28.7 | Parieto-occipital sulcus |
|  | 4 | 389.31 | -28.9 | -84.2 | 12.2 | Occipito-superior and transversal sulcus |
|  | 5 | 330.90 | -49.7 | -9.3 | 46.8 | Precentral gyrus |
|  | 6 | 250.66 | -39.2 | -29.2 | 38.5 | Postcentral gyrus |
|  | 7 | 250.14 | -33.8 | -79.4 | 24.9 | Occipito-superior and transversal sulcus |
|  | 8 | 216.52 | -32.2 | -21.9 | 63.1 | Precentral sulcus |
|  | 9 | 201.87 | -24.0 | -35.7 | 59.7 | Postcentral sulcus |
|  | 10 | 170.11 | -37.3 | -58.0 | 21.9 | Superior temporal sulcus |
|  | 11 | 168.37 | -12.7 | -43.9 | 31.5 | Subparietal sulcus |
|  | 12 | 154.69 | -43.3 | -75.7 | 15.5 | Middle occipital gyrus |
|  | 13 | 142.01 | -33.4 | -3.1 | 43.9 | Precentral sulcus superior part |
|  | 14 | 138.59 | -54.5 | -50.9 | 29.2 | Supramarginal gyrus |
|  | 15 | 120.66 | -46.3 | -55.5 | 8.0 | Superior temporal sulcus |
|  | 16 | 114.03 | -9.6 | -8.4 | 40.4 | Posterior cingulate gyrus and sulcus |
|  | 17 | 104.61 | -11.7 | -94.8 | 11.3 | Superior occipital gyrus |
|  | 18 | 92.79 | -35.4 | -6.0 | 10.6 | Circular sulcus |
|  | 19 | 91.93 | -14.5 | -29.1 | 58.9 | Central sulcus |
| *Right Hemisphere* |  |  |  |  |  |  |
|  | 1 | 2177.40 | 25.9 | -90.0 | -10.9 | Occipital pole |
|  | 2 | 1748.29 | 19.1 | -73.0 | 6.5 | Calcarine sulcus |
|  | 3 | 826.03 | 26.6 | -91.6 | 3.8 | Middle occipital sulcus |
|  | 4 | 721.87 | 44.6 | -64.5 | 25.7 | Superior temporal sulcus |
|  | 5 | 239.42 | 10.5 | -28.8 | 38.3 | pars marginalis of the cingulate sulcus |
|  | 6 | 212.76 | 37.6 | -14.1 | 34.5 | Central sulcus |
|  | 7 | 189.32 | 45.1 | -54.9 | 29.5 | Superior temporal sulcus |
|  | 8 | 166.83 | 36.0 | -79.2 | 31.1 | Middle occipital gyrus |
|  | 9 | 162.42 | 13.4 | -67.6 | 37.8 | Parieto-occipital sulcus |
|  | 10 | 108.84 | 12.4 | -51.3 | 42.1 | Subparietal sulcus |
|  | 11 | 108.61 | 12.2 | -89.9 | 22.3 | Superior occipital gyrus |
|  | 12 | 108.55 | 27.0 | -64.0 | 28.6 | Occipito-superior and transversal sulcus |
|  | 13 | 106.85 | 7.9 | -36.2 | 36.0 | Subparietal sulcus |
|  | 14 | 92.15 | 31.6 | -25.6 | 16.0 | Posterior part of the lateral fissure |
|  | 15 | 88.75 | 32.9 | -55.5 | 39.7 | Intraparietal and transversal parietal sulcus |

